# Terminal Reproductive Investment, Physiological Trade-offs and Pleiotropic Effects: Their effects produce complex immune/reproductive interactions in the cricket *Gryllus texensis*

**DOI:** 10.1101/499236

**Authors:** Atsushi Miyashita, Ting Yat M. Lee, Laura E. McMillan, Russel H. Easy, Shelley A. Adamo

## Abstract

1. Should females increase or decrease reproduction when attacked by pathogens? Two hypotheses provide opposite predictions. Terminal reproductive investment theory predicts an increase in reproduction, but hypothesized physiological trade-offs between reproduction and immune function might be expected to produce a decrease. There is evidence for both hypotheses. What determines the choice between the two responses remains unclear. We examine the effect of age on the reproductive response to immune challenge in long-wing females of the Texas field cricket, *Gryllus texensis*, when fed an ecologically valid (limited) diet.
2. The limited diet reduced reproductive output. However, immune challenge had no effect on their reproductive output either in young or middle-aged crickets, which is contrary to either prediction.
3. Flight muscle maintenance correlated negatively with reproductive output, suggesting a physiological trade-off between flight muscle maintenance and reproduction. Within the long-wing variant there was considerable variability in flight muscle maintenance. This variability may mask physiological trade-offs between immunity and reproduction.
4. Middle-aged crickets had higher total phenoloxidase (PO) activity in their hemolymph, compared to young females, which is contrary to the terminal investment theory. Given that PO is involved in both immunity and reproduction, the increased PO may reflect simultaneous investment in both functions.
5. We identified four proPO transcripts in a published RNA-seq dataset (transcriptome). Three of the proPO genes were expressed either in the fat body or the ovaries (supporting the hypothesis that PO is bifunctional); however, the two organs expressed different subsets. The possible bifunctionality of PO suggests that it may not be an appropriate immune measure for immune/reproductive trade-offs in some species.
6. Increasing age may not cue terminal reproductive investment prior to senescence.

## Introduction

Resources are finite in animals. The partitioning (i.e. allocation) of those resources among resource-intensive traits such as immunity and reproduction can lead to physiological trade-offs, resulting in negative correlations between them (e.g. in insects, Zera & Harshman 2001; Lawniczak *et al.* 2007; Schwenke, Lazzaro & Wolfner 2015; in vertebrates, French et al. 2007, McCallum and Trauth 2007; Kalbe et al. 2009; Nordling et al. 1998; and Mills et al. 2009). Consistent with this argument, activation of an immune response leads to reduced reproductive effort in a range of species (e.g. insects, see Schwenke *et al.* 2016). However, under some conditions, an immune challenge leads to increased reproductive output, which is usually interpreted as a type of fecundity compensation (Duffield *et al.* 2017). An immune challenge signals a risk of early death from infection, and, therefore, a decline in an animal’s residual reproductive value. In response, the animal shifts investment away from somatic maintenance to fuel a final bout of reproduction. This strategy is called terminal reproductive investment (Clutton-Brock 1984). Whether an individual should increase or decrease reproduction when infected depends on a range of poorly understood factors (e.g. age, Duffield *et al.* 2017).

Insects make good model systems for these types of questions, in part because their reproductive output is easy to quantify, and their immune systems are simpler than those of vertebrates (Adamo 2017). Moreover, insects like the cricket *Gryllus texensis* are large enough to measure multiple immune components simultaneously (e.g. key insect immune components such as phenoloxidase (PO) activity (Cerenius, Lee & Söderhäll 2008)).

Age is expected to reduce the fitness benefit gained from increasing immune function and decreasing reproduction (i.e. a physiological trade-off) when infected (see Duffield *et al.*, 2017). Cricket residual reproductive value declines with age (Shoemaker et al. 2006), and, therefore, the fitness pay off for reducing current reproduction to preserve future reproduction should decline over time. Furthermore, if declining immune function due to age (i.e. during senescence) reduces the chance of recovery (e.g. (Adamo, Jensen & Younger 2001)), then it may be adaptive to prioritize reproduction during an immune challenge in older animals. Therefore, age should increase the likelihood that terminal reproductive investment will be activated by an immune challenge (Duffield et al., 2017). Supporting this, interactions between age, immune challenge and reproductive investment have been observed in several male insect models such as *Gryllodes sigillatus* (Duffield *et al.* 2018), *Drosophila nigrospiracula* (Polak & Starmer 1998), and *Allonemobius socius* (Copeland & Fedorka 2012). However, evidence in female insects is relatively scarce, even though reproductive output in females is often easier to quantify. Immune challenge has a variable effect on female crickets (Table 1); possibly female age may help explain this variability. In *G. texensis*, females retain high immunocompetence throughout their adult stage, while in males it declines (Adamo, Jensen & Younger 2001). These results suggest that females may have a reproductive resource allocation strategy that is different from males (also see Rapkin *et al.* (2018) for sex-specific effects of macronutrient intake on trade-offs between reproduction and immunity in *G. sigillatus*). In this study, we examine the effect of age on the female reproductive response to infection.

**Table 1.** Effect of immune challenge on reproduction in female crickets

| Species | Dosage | Age (post adult) at the start of treatment | Duration of immune challenge | Effect on reproduction | Reference |
| --- | --- | --- | --- | --- | --- |
| *Acheta domesticus* | 100 µg/cricket of *Serratia marcescens* LPS<br>5 x 10<sup>4</sup> live cells of *S. marcescens* | 2 weeks<br>2 or 5 weeks | Acute | + (positive)<br>+ | (Adamo 1999) |
| *Acheta domesticus* | 1 or 2 nylon pieces/cricket (implantation) | 18 days | Chronic (3 weeks) | - (negative) | (Bascuñán-García *et al.*, 2009) |
| *Gryllus texensis* | 8.75 x 10<sup>3</sup> live cells of *S. marcescens*<br>1 x 10<sup>5</sup> live cells of *S. marcescens*<br>1.2 x 10<sup>5</sup> live cells of *S. marcescens* | 11 to 19 days<br>11 to 19 days<br>2 weeks | Acute | +<br>0 (no effect)<br>+ | (Shoemaker *et al.* 2006) |
| *Gryllus texensis* | LD<sub>01</sub> of *S. marcescens* live cells<br>LD<sub>01</sub> of *Bacillus cereus* live cells | 2 weeks<br>2 weeks | Acute | 0<br>- | (Adamo & Lovett 2011) |
| *Gryllus texensis* | 20 µg/cricket of *S. marcescens* LPS<br>100 µg/cricket of *S. marcescens* LPS | 13 to 19 days | Chronic (every three days for 12 days) | 0<br>0 | (Shoemaker & Adamo 2007) |
| *Gryllus texensis* | 1 x 10<sup>4</sup> heat-killed *S. marcescens* | 1 day | Chronic (every three days for 17 days) | - | (Stahlschmidt *et al.* 2013) |
| *Hemideina crassidens* | 100 µg *S. marcescens* LPS<br>500 µg *S. marcescens* LPS | Uncontrolled (field collection) | Chronic (every four days for 17 days) | -<br>- | (Kelly 2011) |
| *Gryllodes sigillatus* | Nylon implantation (encapsulation response) | 14 day | Chronic | 0 | Rapkin *et al.* 2018 |

In this study, we chose two age classes to assess the effect of immune challenge in this study: young (11 days as an adult) and middle-aged (21 days old). At 11 days of age (i.e. young crickets), females have mated and begun to produce eggs, but their reproductive activity (oviposition rate) is less than maximal in this cricket (Shoemaker, Parsons & Adamo 2006). By 21 days of age, females are fully mature, with high oviposition rates (Shoemaker *et al.* 2006). However, they are still within the typical age for females found in the field (Murray & Cade 1995), thus they are not old in an ecological sense. We predicted that young female crickets would respond to an immune challenge with decreased reproduction, but an enhancement of immune function (e.g. increased lysozyme-like function). Middle-aged females, on the other hand, would increase oviposition, but would show a more modest activation of their immune response compared with younger females.

One technical difficulty in determining whether there has been a physiological trade-off between immunity and reproduction is assessing immunity. Immune function is made up of multiple components, which can sometimes be traded-off for each other (Adamo 2004b). Immune systems can also reconfigure their molecular network pathways, and therefore a reduction in a single immune component may be mistaken for a reduction in investment, as opposed to a reconfiguration (Adamo *et al.* 2016). Finally, the primacy of different immune pathways can shift depending on the physiological context (Armitage & Boomsma 2010; Piñera *et al.* 2013; Adamo 2014; Adamo *et al.* 2016). Therefore, to monitor immunological investments in crickets, it is important to measure multiple aspects of immune function on each animal. We measured PO, glutathione (GSH, which helps buffer the self-damage caused by PO (Clark, Lu & Strand 2010)), and lysozyme-like activity. PO and lysozyme-like activity respond differently to immune challenges; lysozyme-like activity is inducible in response to pathogen challenge in insects while PO may form a constitutive immune defense (Adamo 2004a).

Although PO activity is commonly used as a proxy for immune function in ecoimmunology, PO is also involved in egg production in insects, which potentially complicates the interpretation of PO levels in female insects. PO is involved in processes such as the tanning of the egg chorion (Li & Christensen 1993; Li 1994) and/or the eggs’ antimicrobial defense (Rizki & Rizki 1990; Abdel-latief & Hilker 2008). In *G. texensis*, PO activity in eggs has also been reported (Stahlschmidt *et al.* 2013). There appears to be several sources of PO in insects, and this may depend on the species. Hemocytes have been viewed as a major source of PO (Cerenius *et al.* 2008; Kanost & Gorman 2008; Lu *et al.* 2014); however, in some insects, the fat body and the ovaries also express POs (e.g. mosquitoes, see Fig. 5 of Cui, Luckhart & Rosenberg 2000). Little is known in insects about how these POs are trafficked between organs, thus it remains unclear whether the hemolymph PO level reflects either immune investment or reproductive investment, or both. To fulfill the knowledge gap on molecular information about PO production in crickets, we assess PO gene expression in both fat body and ovaries.

## Materials and Methods

### Animals

Female *G. texensis* crickets were originally obtained from San Antonio, Texas, USA, and have been maintained in the laboratory for approximately 8 generations. The colony was maintained at 26°C on a 12/12 hour light/dark cycle, supplied with food and water *ad libitum*. Long-winged adult females were weighed and isolated from the colony within 48 hour after the final molt (the day which we call ‘day 1’ in this study). We did not use the short-wing morphs in this study. Each of the isolated females was isolated in a plastic container and supplied with a shelter and water bottle. Food was placed in the individual containers for 3 hours every 3 days. During those 3 hours, crickets could feed *ad libitum*. This diet has been shown to produce females with the same fat content as females collected in the field (Adamo *et al.* 2012). On days 7 and 8 (female adult age), each female was provided with three different males. Each male was placed in the female’s container for about 8 hours. After each mating, males were switched so as to ensure that each female was exposed to three different males. All experiments were approved by the Animal Care Committee of Dalhousie University (# I-11-025) and are in accordance with the Canadian Council on Animal Care.

### Treatments

Female crickets were randomly assorted by weight, and assigned one of the following eight treatments on the day they were isolated from the colony. Days were counted from the day of isolation (see timeline in Fig. 1).

*Early Controls (Control (E))*. Crickets were handled on day 11, and hemolymph samples were collected on day 12 and day 36.
*Late Controls (Control (L))*. Crickets were handled on day 21, and hemolymph samples were collected on day 22 and day 36.
*Early Immune Challenge (IC (E))*. On day 11, crickets were injected with 2 μL of a mixture of heat-killed pathogen cells *(Serratia marcescens, Bacillus cereus* and *Beauveria bassiana*.) The dose of each pathogen was approximately 1/10 of the LD50 dose prior to heat inactivation. Hemolymph samples were collected on day 12 and day 36.
*Late Immune Challenge (IC (L))*. Crickets were injected on day 21 with 2 μL of the same heat-killed pathogen mixture described above. Hemolymph samples were collected on day 22 and day 36.
*Early Sham (Sham (E))*. On day 11, crickets were poked by an empty injection needle, but not injected with any sample. Hemolymph samples were collected on day 12 and 36.
*Late Sham (Sham (L))*. On day 21, crickets were poked by an empty injection needle, but not injected with any sample. The hemolymph samples were collected on day 22 and 36.
*No Treatment Control (NTC)*. Hemolymph samples were collected on day 36.
*No treatment Control (ad lib feeding) (NTC (ad lib))*. Hemolymph samples were collected on day 36 and crickets were fed *ad libitum*.

**Figure 1.**
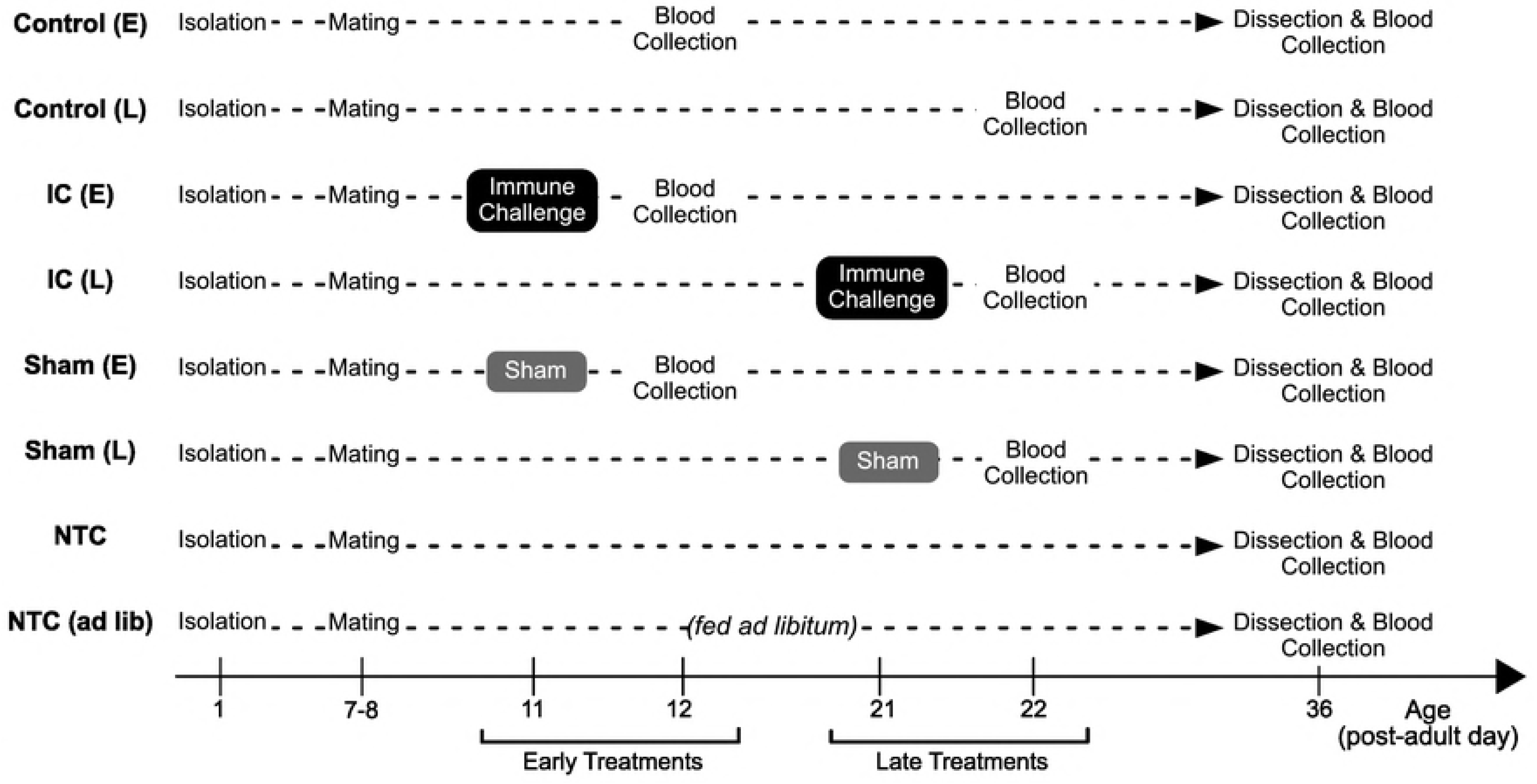
Experimental Schedules. The crickets used in this study (except for ones for the ovarian gene expression experiment) went through one of the eight experimental timelines shown in the chart. Details are described in Materials and Methods.

### Reproductive Output

We monitored the number of eggs laid in the cotton balls twice a week. Five eggs were subsampled from each cotton ball and placed separately in centrifuge tubes (1.5mL) with a small piece of cotton and 500 μL of water. In cases where the number of eggs laid in the cotton ball was less than five, all eggs were sampled. These eggs were then kept at 26°C and monitored for 35 days. Hatch date, hatchling survival (daily), and hatchling body mass at 35 days after the hatch day were monitored for the sampled eggs. Eggs were censored and assumed not viable if they had not hatched within 35 days. The reproductive value (RV), a proxy for fitness, was calculated as a product of the number of eggs and the hatch ratio. For example, if a female laid 50 eggs and 3 out of the 5 subsampled eggs hatched, then the reproductive value would be 30.

### Dissection

If still viable, crickets were dissected on day 36. For each cricket, we: 1) measured body mass, 2) collected fat body tissues for gene expression analyses (described below), 3) counted eggs in the lateral oviducts, and 4) observed flight muscle state (functional/histolysed as described by Zera 2003). We also measured length of the hind leg femurs and recorded the average of the two legs. For gene expression analysis in the ovaries, we dissected a group of females on day 15 that were independent of the rest of the study and collected both the fat body and the ovaries. These crickets were also given the intermittent diet during adulthood.

### Hemolymph Collection

We collected hemolymph samples by poking the membrane under the pronotum plate with an ice-cold pipette tip (to retard coagulation), and the hemolymph was collected as it exited the wound. We collected 8 μL of hemolymph which was mixed with 55 μL of ice-cold MilliQ water in a 1.5 mL centrifuge tube. Samples were then split into three fractions (20 μL for the PO and Bradford assays, 23 μL for the GSH assay, and the rest (20μL) for the Lysozyme assay). Immediately after the sample collection, we spun the hemolymph sample for the GSH assay (23 μL) at 18,800 g for 10 min at 4 °C, and 20 μL of the supernatant was immediately mixed with 20 μL of 100 mg/mL meta-phosphoric acid. After 5-minute incubation at room temperature, we spun the samples at 2,900 g for 3 min at room temperature. 35 μL of the deproteinated supernatant was collected in a new 1.5 mL centrifuge tube. All the samples for PO, Bradford, GSH, or Lysozyme assays were stored at −80°C until use.

### Hemolymph Assays

Total PO activity and total protein concentration were measured as described previously (Adamo 2004a). GSH concentration was measured as described previously (McMillan, Miller & Adamo 2017). A detailed information for the assays is described in the Supplementary file. Briefly, for lysozyme-like activity, hemolymph samples were collected as described above. Samples were thawed and spun at 12,000 g for 3 min at 4 °C. 5 μL of the supernatant was mixed with 45 μL *Micrococcus luteus* cell (Sigma #M3770) suspension (10 mg/20 mL Phosphate-buffered Saline (PBS), pH = 7) in a 96-well (flat bottom) plate. The mixture was incubated at 30 °C, and we measured OD_450_ every 30 seconds for 50 minutes. Lysozyme derived from chicken egg white was used to produce a standard curve (Sigma-Aldrich, #62971-10G-F). A blank (PBS) were run concurrently. The mean value from triplicate technical replicates was used for each sample.

### Gene Identification

To identify the gene transcripts in the cricket, we first constructed a transcriptome database based on a raw sequence of RNA reads available online (National Center for Biotechnology Information (NCBI, https://www.ncbi.nlm.nih.gov/). The accession number of the bioproject is PRJNA429132 (submitted by Natural History Museum, Berlin, Germany). We then set up a searching pipeline (written in Python programming language, the code is available at the author’s GitHub repository at https://github.com/atmiyashita/CricketGeneFinder2018/). The code: 1) fetches cDNA sequences in arthropods from NCBI Nucleotide database that are associated with the target protein name (i.e. ‘vitellogenin’, ‘phenoloxidase’ etc.), 2) runs BLAST locally using the fetched sequence as a query and the transcriptome (of *G. texensis*) as a database, 3) outputs the result in xml format, and 4) returns a summary. The hit sequences were then confirmed by blastx at https://blast.ncbi.nlm.nih.gov/Blast.cgi?PROGRAM=blastx&PAGE_TYPE=BlastSearch&LINK_LOC=blasthome to confirm its homology at amino acid sequence level (i.e. primary structure). For vitellogenin, we further performed a physiological validation in this study, because the sequence similarity was relatively low (compared to proPO, see figure S1).

### Gene Expression Analysis

The primers used in this study are listed in Table S1. We followed the MIQE guideline (Bustin *et al.* 2009; Taylor *et al.* 2010) for the qPCR experiments. The fat body (the speckled white tissues found in the abdominal cavity) was collected carefully to minimize collecting other tissues such as the trachea. The ovaries were collected carefully so as not to contaminate the sample with fat body. We washed ovaries once with PBS to further avoid potential contamination of the sample with hemocytes. The tissues were stored in 300 μL of RNAlater (Thermo Fisher Scientific, #AM7020) in 1.5mL centrifuge tubes and frozen at −80°C until further use. Detailed information for RNA extraction, cDNA synthesis, and quantification is described in the Supplementary information.

### Data Analysis

In this study, we isolated 240 female adult crickets assigned across 8 treatment groups. 8 out of the 240 crickets did not mate (i.e. 232 crickets contained spermatheca filled with sperm when dissected). These 8 were excluded from the analysis. 66 crickets were also excluded from the analysis because some data were lost (e.g. due to death, Fig. S2). Thus, we acquired a complete dataset on 166 female crickets that: 1) mated, 2) survived for 36 days. We noted some inter-trial variation in the baseline level of immune factors, so we treated the trial numbers as a random factor in the linear mixed models. Unless stated otherwise, we used packages ‘lme4’ and ‘lmerTest’ running on R (version ‘Short Summer’ (3.4.2)) for the linear mixed models. The details for each analysis is described in the figure legends. As a measurement of condition, the Scaled Mass Index (SMI) (Kelly, Tawes & Worthington 2014) was calculated for each cricket.

## Results

### Effect of food availability on overall reproductive output

Controls that were food-limited (NTC) produced fewer eggs than controls fed *ad libitum*. (i.e. (NTC (*ad lib*)) (Fig. 2A). *Ad lib* fed crickets also laid more eggs, and had more eggs in the lateral oviducts on day 36 than did food-limited crickets (Fig. 2B and 2C). The ratio of eggs laid by day 36 over total eggs produced, was similar between the two feeding conditions (Fig. 2D). The relative number of eggs laid compared with eggs held in reserve in the lateral oviduct did not vary with diet (Student’s t-test; t = 0.87, df = 29, p = 0.39). Also, 3 out of 20 females had pink (functional) flight muscles on day 36 in NTC group, compared to 3 out of 19 females in NTC (*ad lib*) group, suggesting that the feeding condition did not affect the likelihood of flight muscle histolysis by day 36 (Chi-squared test; χ^2^ = 1.3e-31, df = 1, p = 1).

**Figure 2.**
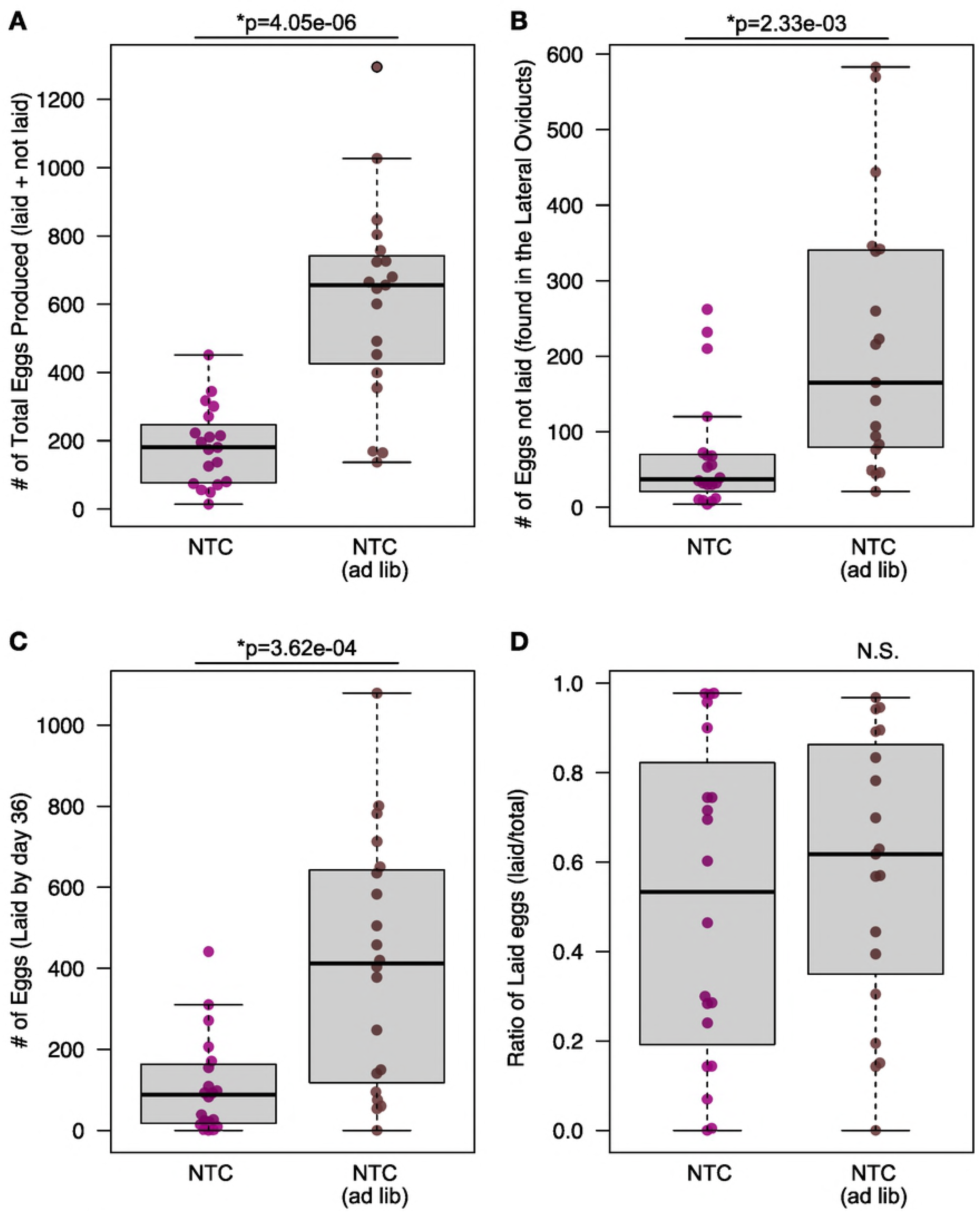
Effect of feeding condition on reproductive output. The feeding condition (intermittent vs. *ad libitum*) significantly affected the reproductive output of the crickets. (A) The total number of eggs produced by day 36 (a sum of the laid eggs and the eggs found in the lateral oviducts) was significantly lower in the NTC than NTC (*ad lib*). (B,C) The number of eggs laid by day 36 (B) and the number of eggs found in the lateral oviducts (C) were also significantly lower in the intermittent feeding condition. The p-values shown in the figure is calculated by Welch Two Sample t-test. (D) The portion of laid eggs (i.e. laid:total ratio) was comparable between the two feeding conditions.

### Treatment effects on immune function and survival

IC (E) and IC (L) groups did not show any differences in PO level, GSH level, and PO:GSH ratio in the hemolymph 24 hours after the immune challenge (i.e. day 12 and 22) compared with controls (Fig. S4). The hemolymph protein concentration in the IC (E) group increased relative to controls 24 h after the immune challenge (Fig. S4). IC (L) showed higher lysozyme-like activity relative to controls 24 hours after the challenge (Fig. S4). Only one group produced effects that lasted until day 36, the IC (L) on GSH (Fig. S4). The overall survival rate on day 12, 22, and 36 was 100% (210/210), 99% (207/210), and 79% (166/210) as shown in Fig. S2, and there was no difference in survival across the seven groups (Fig. S2B; χ^2^ = 9.8, df = 6, p = 0.13).

### No effect of immune challenges on reproductive output

The total number of eggs produced, the number of eggs laid, and the number of eggs in the lateral oviducts, and the ratio of laid eggs to the total number of eggs on day 36 was comparable across all seven treatment groups (Fig. 3). No effect of treatment was observed at any time point (Fig. S5).

**Figure 3.**
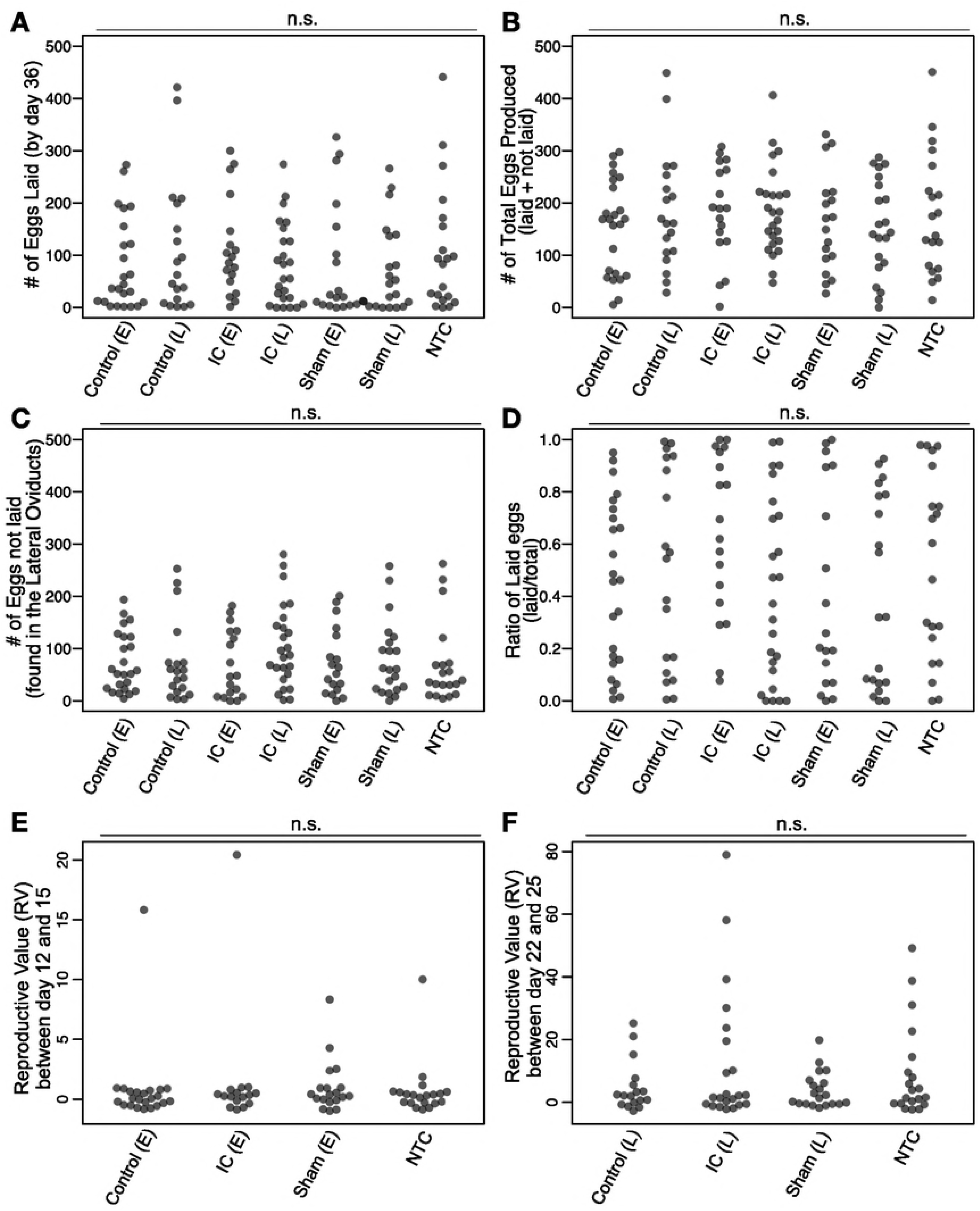
No effect of immune challenge on reproductive output. There was no significant effect of treatments on overall (36-day) reproductive outputs measured as (A) number of eggs laid, (B) number of total eggs produced, (C) number of eggs found in the lateral oviducts, and (D) ratio of laid:total number of eggs. Also, there was no acute effect on the reproductive output immediately after the treatments (E and F). Detailed time course is also shown in Fig. S5, which corroborates the lack of treatment effect on reproductive output.

### Association between dispersion capability and reproductive output

Most crickets had white (histolysed) flight muscle on day 36 (130 of 150 observations), which was observed equally across the eight groups (χ^2^ = 3.9, df = 7, p = 0.79). The crickets that still retained pink (functional) flight muscles on day 36 produced and laid fewer eggs than the crickets with histolysed flight muscle (Fig. 4A, B). The number of eggs found in the lateral oviduct on day 36 was not significantly different between the two morphs (Fig. 4C), but the laid:total ratio was significantly lower in pink-muscle crickets (Fig. 4D). The body condition measure, SMI, was comparable between the two morphs on day 1, but was higher in the white-muscle morph on day 36 (Fig. 4E). Also, an increase in SMI over time was observed in the white-muscle morph, but not in the pink-muscle morph (Fig. 4E). The body mass on day 1 was significantly higher in pink-muscle crickets, but that on day 36 was comparable between the two morphs (Fig. 4F). Increase in body mass over time was only observed in the white-muscle morph (Fig. 4F).

**Figure 4.**
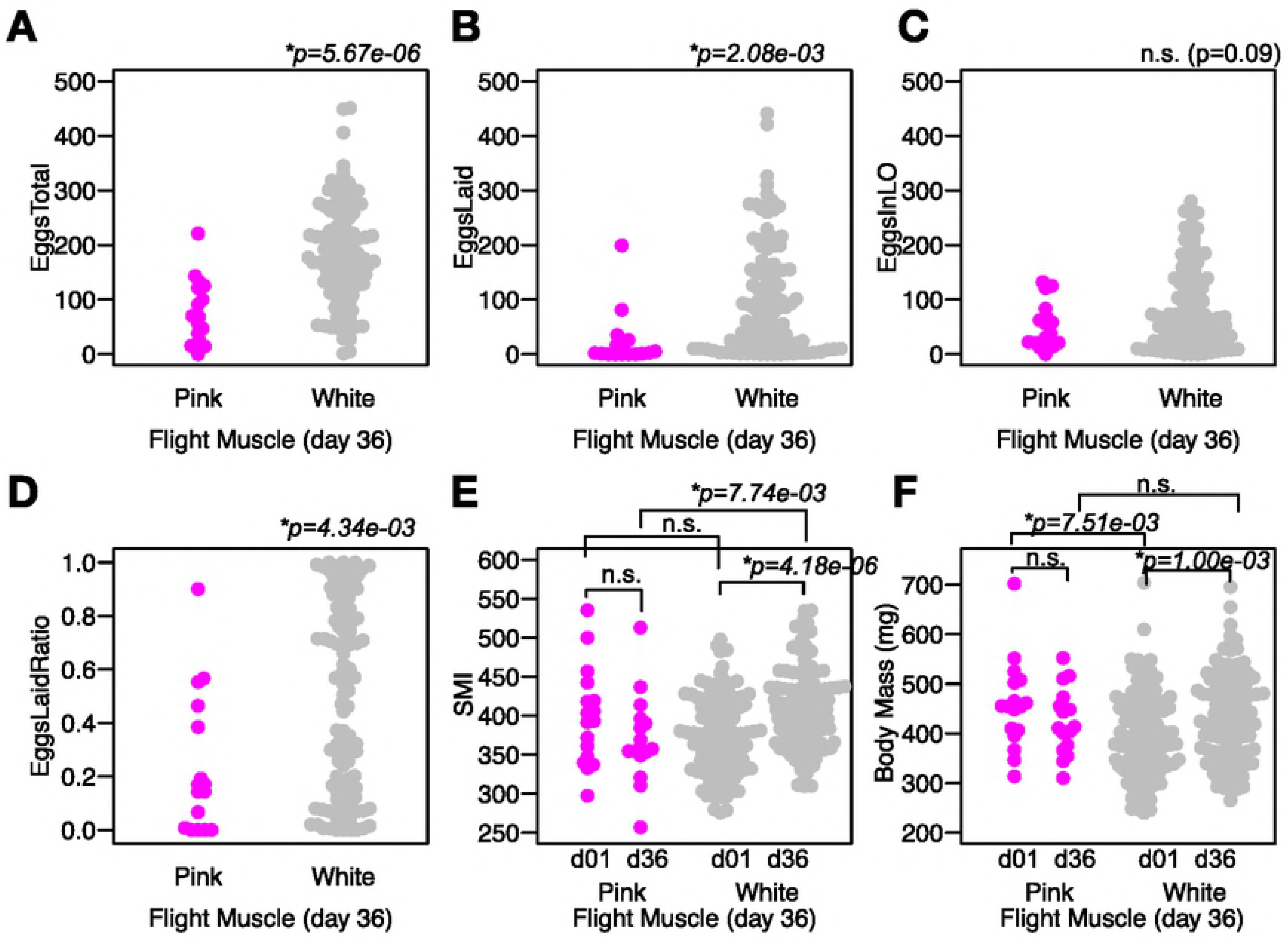
Flight capability is negatively correlated with reproductive output. Total number of produced eggs (A), number of laid eggs (B), and the laid:total ratio (D) was higher in the crickets that had histolysed (white) flight muscle on day 36. The number of eggs found in the lateral oviducts showed a trend toward significance (C). SMI represents Scaled Mass Index (E). d01 and d36 represents day 1 and day 36 (E and F).

### Age-dependent changes in immune measures

Phenoloxidase activity was higher in 22 day old and 36 day old adult females than in 12 day old adult females (Fig. 5A), Other parameters we measured in the hemolymph (GSH, PO:GSH ratio, lysozyme-like activity, and protein level) did not show age-dependent differences (Fig. 5B-E). The PO level showed a modest positive correlation with reproductive output (r-squared = 0.11, p=1.03e-07) (Fig. S6).

**Figure 5.**
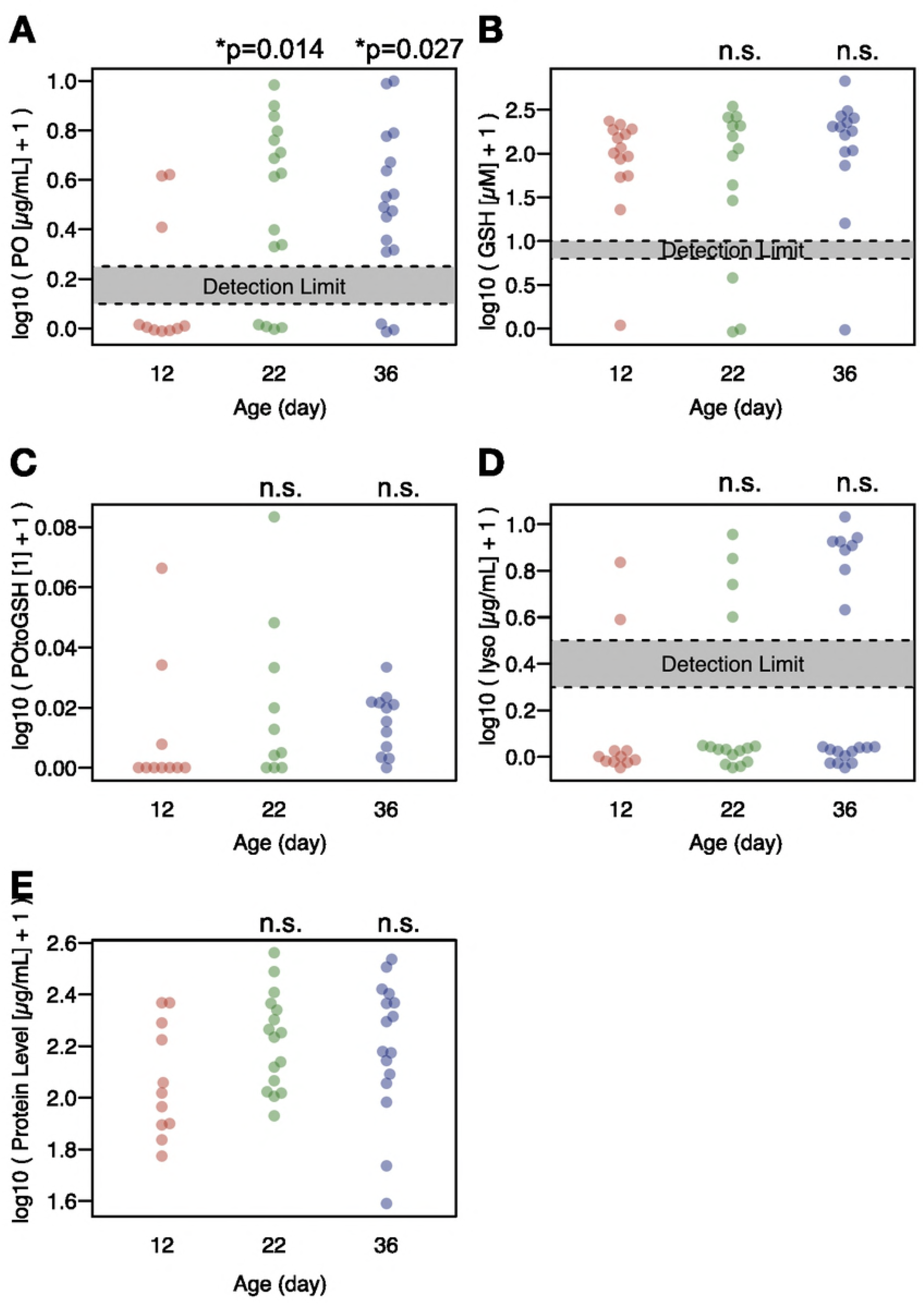
Age-dependent increase of PO level. Age-dependent change in hemolymph parameters. The PO level (μg (tyrosinase equivalent)/mL), the GSH level (μM), the PO:GSH ratio, the lysozyme-like activity (μg/mL), and the protein level (μg/mL) are log10-transformed and shown in the chart. The numbers below each chart represent the ages at which the blood was collected. For days 12, 22, and 36, results from Control (E/L) and NTC are plotted. Each dot represents an individual cricket. The treatment effects were examined using Mixed Linear Models (using ‘lme4’ package in R), considering cohort identity (i.e. experimental date) as a random factor in the model. Only PO showed a significant increase over age (A), while other parameters showed no trend (B-E).

### Expression of proPOs in the ovaries

We detected three proPO transcripts in the ovary and the fat body. proPO1 was consistently expressed in the fat body and the ovaries at comparable levels (Fig. 6A). proPO2 was expressed specifically in the ovaries (Fig. 6B), while proPO3 was expressed specifically in the fat body (Fig. 6C). Vitellogenin was expressed specifically in the fat body (Fig. 6D).

**Figure 6.**
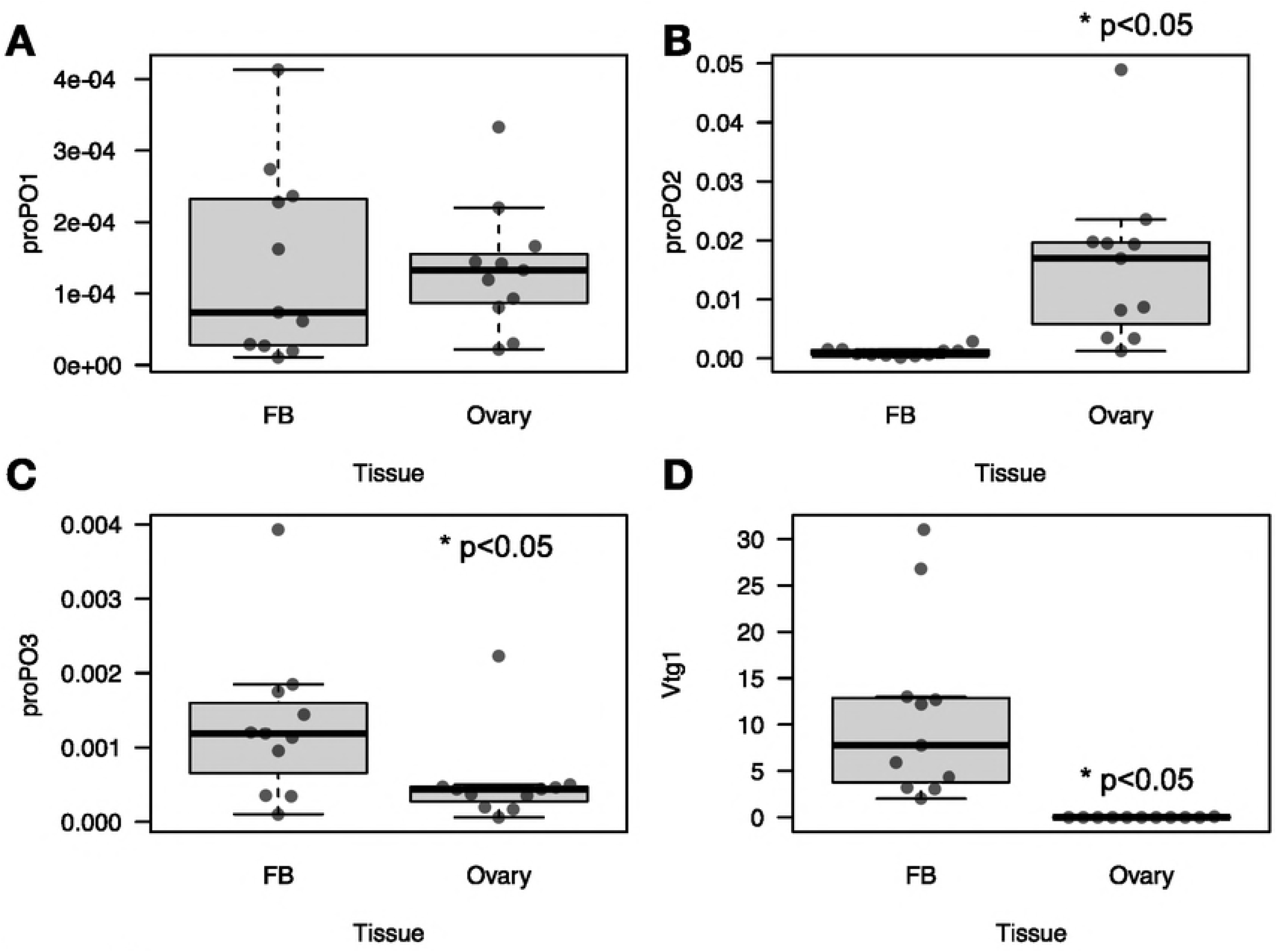
Ovaries produce phenoloxidase (PO) in the cricket. Gene expression levels of proPOs (A-C) and Vitellogenin (D) were measured in the fat body and the ovaries. Experimental procedures are described in Materials and Methods. Translated sequences of each transcript is shown in Figure S1. The values were normalized by two reference genes, and the relative expression levels (arbitrary units), where the expression level for the reference is set to be 1.0.

## Discussion

Food limitation reduced the total number of eggs produced, corroborating earlier studies that this diet reduces reproduction (e.g. Adamo *et al.* 2012). We had assumed that under low food availability, females would have proportionately fewer eggs in reserve in the lateral oviducts than did *ad lib* controls. In other words, we expected that food-limited females would lay proportionately more of their eggs in order to maintain egg output even as egg production fell. However, females maintained the same proportion of eggs in reserve when food-limited as when resources were abundant. Possibly females reduce their risk of low offspring survival by laying eggs in different places at different times. This strategy of oviposition site diversification would explain why females are found with eggs in their lateral oviducts even in the field (Adamo 1999).

Despite the evidence that the food-limited diet reduced the resources needed for reproduction (i.e. because food-limited crickets had fewer total eggs), we found no evidence of a physiological trade-off between immunity and reproduction in either young or middle-aged adult females in response to an immune challenge. This result corroborates other studies in crickets that did not find a reduction in reproduction after a single immune challenge or repeated immune challenges (Table 1). Previous studies on this species have demonstrated that the immune challenge we used induces a robust immune response in female crickets (Adamo 2004a; 2010). Moreover, in this study, we found an increase in lysozyme-like activity 24 h later in older (middle-aged) females, suggesting that the minimal effect was not due to a lack of immune response. However, it is possible that the heat-killed challenge did not induce an enough effect to trigger a physiological trade-off, although the immune challenge does trigger sickness behaviours in this cricket (Sullivan, Fairn & Adamo 2016). It is also possible that the decrease in reproduction was small, which was not noticeable given the large variability in egg number and egg-laying behaviour. Assuming the same effect size and variability as found in our data set (for example, the effect size (Cohen’s d) was 0.05 between the numbers of eggs laid in NTC and IC (E) groups), we would need more than 780 crickets/group to potentially find a positive effect. Such a small effect is inconsistent with most studies on immune/reproductive trade-offs (e.g. Stahlschmidt *et al.* 2013).

This discrepancy may be explained by unique aspects of cricket life history. Long-winged *G. texensis* crickets histolyze their wing muscles at some point during their adult life (authors’ personal observation), which releases additional resources for reproduction in other cricket species (Zera, Sall & Grudzinski 1997; Zera, Potts & Kobus 1998). Once the muscles are histolyzed, they are no longer capable of flight (Zera & Denno 1997). The additional resources provided by the wing muscles may reduce trade-offs between immunity and reproduction. Mathematical models of trade-offs have demonstrated that trade-offs may be difficult to demonstrate if the variance in resource acquisition is large compared with that in resource allocation (van Noordwijk & de Jong 1986; Zera & Harshman 2001; Metcalf 2016). In this study we observed large variability in the timing of flight muscle histolysis within each group, suggesting that there is considerable individual difference in available resources at any particular time point. Supporting our hypothesis that flight muscle histolysis may provide an important boost in resources for reproduction, flight muscle histolysis showed a strong association with reproductive output. Future studies should note whether wing muscles have been histolyzed when studying physiological trade-offs in crickets.

There was also no evidence of terminal reproductive investment in either age class. We expected that older (i.e. middle-aged) female crickets should increase reproduction when given heat-killed bacteria, as has been observed previously (Shoemaker *et al.* 2006). However, there was no evidence that females became more sensitive to an immune challenge with age. In previous reports from our laboratory, female crickets showed terminal reproductive investment, even at a young age (Adamo 1999; Shoemaker *et al.* 2006), but those immune challenges were close to a lethal dose. The effect of a sub-lethal dose of bacteria on reproduction was only observed in *G. texensis* when moist sand was used for egg-laying substrate, and was not observed when moist-cotton was used, as in this study (Shoemaker *et al.* 2006). *G. texensis* females prefer to oviposit in moist sand over moist cotton (Shoemaker *et al.* 2006), which may have affected the terminal investment thresholds. This point needs to be validated in future studies.

There is an alternative explanation for the lack of terminal investment with age in this study: the threshold for terminal reproductive investment may not decrease with age until the beginning of senescence. Although crickets can survive for more than 8 weeks in the laboratory, they show signs of senescence after only 4 weeks (Shoemaker *et al.* 2006). This is consistent with our study that showed an increase in mortality only at the last time point (i.e. 36 days, Fig. S2). Crickets were not given an immune challenge at this time point, and, therefore, were not tested for terminal reproductive investment at a time when mortality due to age was increasing. Prior to senescence, the risk of death for female crickets may be the same each day, unless predator or pathogen prevalence increases. Decreasing the threshold for terminal reproductive investment prior to senescence may not be advantageous for female crickets when oviposition site is important for offspring survival (Shoemaker et al., 2006) and optimal oviposition sites are not available.

Instead, females may depress egg production and/or egg laying, even when infected, until conditions are favourable for offspring development. In some females of this species, completing a dispersal flight may also signal better oviposition opportunities. Flight is known to increase egg production in this and other cricket species (Guerra & Pollack 2009; Zeng, Zhu & Zhao 2014), although some long-winged females histolyze their flight muscles at a young age even without a dispersal flight (Zera, Sall & Grudzinski 1997). The threshold for terminal reproductive investment may be very high prior to wing muscle histolysis. Given that this event occurs at different dates across individuals, the effect may be masked by the number of non-responders.

Contrary to the terminal investment theory, in which individuals are assumed to invest less in somatic maintenance with age, we found an age-dependent increase in PO. This is consistent with an earlier study that found an age-dependent increase in PO in female, but not in male, *G. texensis* (Adamo *et al.* 2001). An increase in PO activity with age has been found in females of other species, and this increase can lead to Malpighian tubule damage in old female *Tenebrio molitor* (Khan, Agashe & Rolff, 2017). Whether PO-induced damage is involved with the increase in mortality observed by us on day 36 is unknown. However, given that the PO:GSH ratio remained constant across ages (GSH buffers the self-damaging toxicity of PO), the self-damaging cost of the increased PO may be minor.

The increase in PO activity with age may not represent an increase in immune investment, or be an example of immune dysregulation. The increase in PO may represent an increase in reproductive investment. PO is needed for the tanning and defense of insect eggs (Rizki & Rizki 1990; Li & Christensen 1993; Li 1994; Abdel-latief & Hilker 2008), and in some insects, it appears to be synthesized by sources outside of the ovary and transported to the ovaries through the blood (e.g. mosquito (Kim *et al.* 2005)). Therefore, increases in PO hemolymph levels may represent an increase in reproductive effort, which incidentally also increases the amount of PO available for immunity, and may lead to immunopathology in old age (Khan, Agashe & Rolff 2017). We have 3 lines of evidence for this in our study. PO activity rises in middle-aged (day-22) crickets, which is prior to senescence (Fig. S2). Middle-aged females have a high reproductive output, consistent with an increased need for PO to maintain increasing egg production (Fig. S5). The age-dependent increase in PO is observed only in female *G. texensis* (Adamo *et al.* 2001), suggesting that the rise of PO is involved with female-specific life-history traits such as egg production. Finally, we detected proPO gene expression in the fat body, and, therefore, it could supply PO to both hemolymph and ovary. However, we also found that the ovary expressed two subtypes of proPO genes, and, therefore it is uncertain to what extent the ovary and fat body contribute to egg PO. We have not done an in-depth molecular analysis of the proPOs expressed in the ovaries, but the differential expression of proPO subtypes between the fat body (primarily an immune organ, but also involved in reproduction via vitellogenin (yolk protein) production, Arrese & Soulages 2010)), and the ovary indicates the complexity and pleiotropic nature of PO. Identifying the circulating PO subtype(s) in the hemolymph and in the eggs would help answer this question. It is these types of mechanistic details that are needed to understand trade-offs (e.g. Zera and Harshman 2001; Duffield *et al.* 2017). This complexity also suggests that PO is not an ideal proxy for immune investment in immune/reproductive trade-off studies in female insects.

The lack of effect of age on the terminal reproductive response to infection in female crickets in this study reflects the generally weak effect of age on the reproductive response to infection in other female animals (Duffield et al., 2017; Adamo 1999; Sanz et al. 2001; Cotter et al. 2011). These results suggest that age has little effect on the threshold for terminal reproductive investment in females during an infection prior to senescence.

## Authors’ Contribution

AM and SA conceived the ideas and designed methodology; RE established the qPCR methods; AM, ML, LM collected the data; AM and SA analyzed the data; AM wrote R and Python codes; AM and SA led the writing of the manuscript.

## Acknowledgment

We thank Beatrice Chiang, Lynn Ann and Lindsey Puddicombe for cricket colony maintenance. We also thank Anne Johnsen, Madeline Fawley, Brieanna Limkilde, Bakhmala Khan and Galen Pickett for help with data collection. We thank Ian Weaver for providing equipment. This study was supported by an JSPS (Japan Society for the Promotion of Science) Overseas Research Fellowship to AM and an NSERC (Natural Science and Engineering Research Council of Canada) Discovery Grant to SA.

